# Integrative Omics for Informed Drug Repurposing: Targeting CNS Disorders

**DOI:** 10.1101/2020.04.24.060392

**Authors:** Rammohan Shukla, Nicholas D Henkel, Khaled Alganem, Abdul-rizaq Hamoud, James Reigle, Rawan S Alnafisah, Hunter M Eby, Ali S Imami, Justin Creeden, Scott A Miruzzi, Jaroslaw Meller, Robert E. Mccullumsmith

## Abstract

The treatment of CNS disorders, and in particular psychiatric illnesses, lacks disease-altering therapeutics for many conditions. This is likely due to regulatory challenges involving the high cost and slow-pace of drug development for CNS disorders as well as due to limited understanding of disease causality. Repurposing drugs for new indications have lower cost and shorter development timeline compared to that of de novo drug development. Historically, empirical drug-repurposing is a standard practice in psychiatry; however, recent advances in characterizing molecules with their structural and transcriptomic signatures along with ensemble of data analysis approaches, provides informed and cost-effective repurposing strategies that ameliorate the regulatory challenges. In addition, the potential to incorporate ontological approaches along with signature-based repurposing techniques addresses the various knowledge-based challenges associated with CNS drug development. In this review we primarily discuss signature-based *in silico* approaches to drug repurposing, and its integration with data science platforms for evidence-based drug repurposing. We contrast various *in silico* and empirical approaches and discuss possible avenues to improve the clinical relevance. These concepts provide a promising new translational avenue for developing new therapies for difficult to treat disorders, and offer the possibility of connecting drug discovery platforms and big data analytics with personalized disease signatures.

## I. Introduction

Central nervous system (CNS) disorders are the leading cause of disability worldwide. There is a substantial amount of research dedicated to understanding the molecular, cellular, and neurobiological underpinnings of CNS disorders, but integration of these lines of inquiry for drug repurposing is lacking. The present challenges for drug discovery for brain disorders, as summarized by the Institute Of Medicine’s Forum On Neuroscience And Nervous System Disorders, can be broadly categorized into two domains: regulatory affairs and knowledge discovery [1]. At the regulatory domain, the failure of late-stage clinical trials for neuropsychiatric disorders is disproportionately high. According to the Food and Drug Administration (2013), only 2 out of 27 (7%) recently approved drugs had indications for CNS disorders [1,2]. Overall drug development for CNS disorders is associated with high risk of failure, inordinate time to reach the market, and financial cost greater than the motivation to treat a large population in need. In effect, large pharmaceutical companies are led to withdraw investments from the development of therapeutics for CNS disorders. This regulatory challenge is tightly coupled to the second barrier to drug discovery, the knowledge domain. Amidst the complexity of the brain, the pathogenesis and etiology of CNS disorders is not well-characterized: there is a lack of biomarkers, molecular targets, and appropriate models, whether it be *in vivo* or *in vitro*, to fully recapitulate the disorders in question [3]. While we wait for the promise of modern neuroscience, genome-based technologies and large consortia of “omics” data provide an opportunity to strengthen therapeutic options for CNS disorders, and particularly for psychiatric diseases, through drug repurposing approaches.

Drug repurposing is the practice of identifying a known drug and redeveloping it for a new indication (Table 1). As the repurposed drug has already been approved by the Food and Drug Administration (FDA), the economic burden of testing for tolerability, safety, side effects, efficacy, and the regulatory affairs associated with phase I trials is diminished, thereby making it a cost-effective approach. While a new therapeutic would need extensive characterization of the aforementioned parameters, the repurposed drug would only need to be characterized for the new indication of interest [4,5]. Combining drug repurposing with more sophisticated characterization of receptors, pathways, and effectors in the CNS provides an efficient opportunity to identify unexplored disease mechanisms and address the challenges associated with the knowledge domain shortfall described by the Institute of Medicine [1].

**Figure 1.**
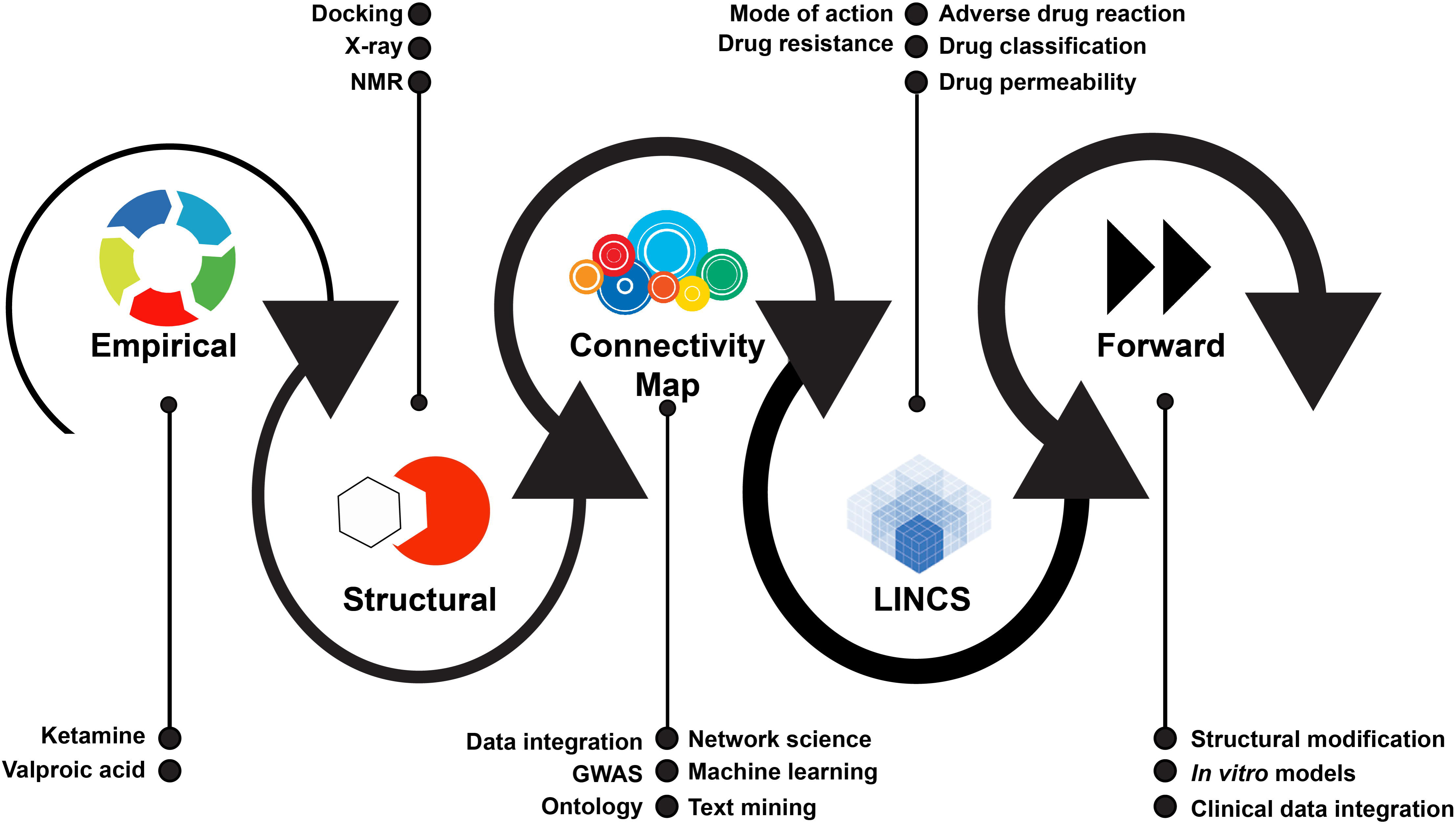
Chronology of drug repurposing approaches: The thickness of each circle depicts the relative expansion of each approach in terms of the number of structure or signature profiles available and potential to identify new candidate. Various data integration approaches can be used to analyze structure and signature based data to identify new therapeutic indication and knowledge mining efforts.

**Table 1.** Glossary of relevant bioinformatic terms

| **Term** | **Definition** |
| --- | --- |
| Drug Repurposing | Repurposing (also called repositioning, reprofiling, retasking) is the investigation and application of existing drugs for a new therapeutic purpose |
| Connectivity | A measure of match or mismatch between two signatures emerging from different sources (drug, disease or pathway). A match defines a positive connection where a mismatch defines a negative connection. Occasionally, similarity, concordance and correlations are used in similar context. |
| Filtering | A user selected subset of data based on chosen search characteristics |
| L1000 | A fluorescence-based assay to measure the gene expression levels of 978 transcripts, which are highly representative of the transcriptome. The 978 representative genes are called the landmark genes |
| LINCS | Library of Integrated Network-Based Cellular Signatures. A repository of signatures from diverse cell types against variety of perturbing agent. |
| Perturbagen | Substance used to treat the cells in order to generate signature associated with it. Includes shRNAs, cDNAs, small molecules, drugs and biologics. |
| Query | A user defined list of gene for which the connectivity score is calculated with respect to all the profiles available in a database. |
| Signature | A characteristic pattern of gene expression associated with a phenotype, response to disease or cellular response to drug perturbation. |

There is promise for new pharmacologic therapies to be implemented for a number of CNS disorders and the source of these new therapies may already be present in the field [6]. For example, within the realm of psychiatric medicine, a recent study suggested that drugs originally designed to treat a psychiatric disorder may be repurposed for other CNS disorders. 77% of source drugs (i.e. new drugs approved by the FDA) originally indicated for a psychiatric disorder have been repurposed three or more times [6]. Repurposing for Alzheimer’s disease has occurred the most, followed by substance use disorders, bipolar disorder, depression, neuropathy, multiple sclerosis, and schizophrenia. Such repurposing efforts have been largely empirical (Table 2), if not accidental, suggesting a single drug has multiple targets and can crossover into different systems for seemingly disparate conditions. This promiscuity has been exploited to bolster drug-repurposing efforts in psychiatry.

**Table 2.**
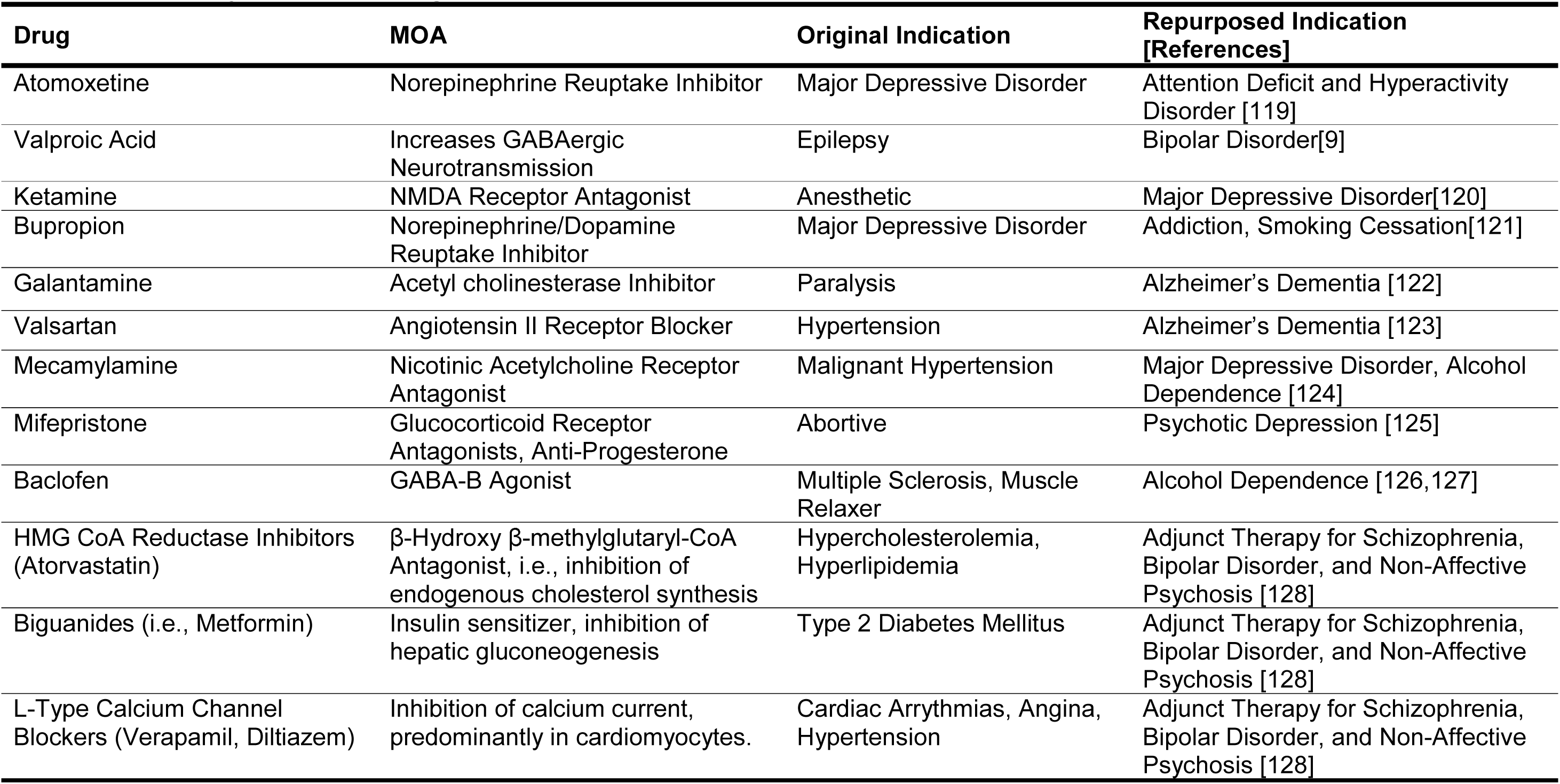
Empirically repurposed drugs for CNS disorders

### Examples of empiric drug repurposing

The most successful examples of empiric drug repurposing in psychiatry include the utilization of valproic acid and ketamine (Table 2). Valproic acid has anticonvulsant properties and shortly after its initial characterization for epilepsy, a therapeutic effect was established for bipolar disorder. The FDA approved divalproex sodium, a modified valproic acid, for the mania phase of bipolar disorder [7,8]. Valproic acid is considered more effective than lithium and more tolerable than medications such as carbamazepine or olanzapine, which are also indicated for bipolar disorder [9]. Besides its repurposing for bipolar disorder, valproic acid is still commonly used as an anti-epileptic [10].

Ketamine, a dissociative anesthetic, was recently repurposed for its anti-depressant effects [11]. 20% of individuals who receive some type of pharmacological therapy for depression do not respond, rendering them treatment resistant [12,13]. Various open label case studies and randomized controlled trails have indicated that ketamine serves as an effective therapeutic for treatment resistant depression [14]. While major depressive disorder is largely considered a disorder of monoaminergic dysregulation, ketamine is an NMDA-receptor antagonist and its repurposed use has led to the characterization of glutamatergic dysregulation in depression, an example of knowledge discovery following empirical drug repurposing.

### Examples of bioinformatics based drug repurposing

In the past five years, with the advent of omics technologies, more sophisticated methods have been used to identify drugs for repurposing in psychiatric medicine [15]. One promising approach involves transcriptional profiling of cell lines after pharmacologic exposure, generating a drug signature that provides mechanistic clues for the downstream effects of the drug. A repository of such signatures can be readily used to repurpose drugs for new indications. Two notable examples using this approach pertaining to depression [16] and Alzheimer’s dementia (AD) [17] are discussed below.

With the goal of identifying new MDD treatments, human hippocampal neural progenitor cells were treated with escitalopram, a selective-serotonin reuptake inhibitor, or nortriptyline, a tricyclic antidepressant [16]. Changes in cellular transcriptomes were characterized and used to develop an “antidepressant” transcriptional signature [18,19]. Using transcriptomic based drug-repurposing approaches, the following candidates were identified: Clomipramine (a tricyclic antidepressant), W-7 (an intracellular calmodulin antagonist), and vorionstat (a histone deacetylase inhibitor). One of these drug candidates confirms this methodology, while the other two provide possible new leads and mechanisms of action that may be explored for the treatment of MDD.

Following mining of the Gene Expression Omnibus (GEO) database for transcriptomic signatures of AD from postmortem brain and animal models, bioinformatics approaches were used to identify compounds with therapeutic potential [17]. Treatment of induced pluripotent stem cells differentiated into adult cortical neurons with candidate compounds from these bioinformatics analyses led to identification of pathways related to bioenergetics. These results support a role for impaired mitochondria in the pathogenesis of AD, yielding candidate drugs that may be exploited as a therapeutic agent for AD [20,21].

The past success of drug repurposing in psychiatry, both through empiric observation (Table 2) and bioinformatics approaches, encourages greater investment in high-throughput, systematic approaches for drug-discovery. Transcriptomic approaches, for instance, can be implemented with increasing ease due to their low cost, public availability of datasets, and ability to incorporate advanced network and systems biology approaches for causal ontology-based drug repurposing. Thus, in this review we emphasize transcriptional profiling as a major new development for drug repurposing efforts. We also provide an overview of different approaches, resources and data integration methods utilized to repurpose drugs that may be particularly relevant for psychiatric disorders. Present limitations and other potential avenues where drug-repurposing approaches can be deployed will also be discussed.

## II. Approaches for Drug Repurposing

Omics data can be used for systematic *in silico* repurposing through structure- and/or signature-based methodologies. Structure based methods take advantage of resolved (or modeled) 3D structures of relevant protein targets and aim to identify potential drug-like modulators by assessing shape complementarity and strength of binding between the target and ligands [22]. Signature-based methods use the transcriptomic “fingerprints” of disease states and animal models, as well as the effects of drugs on *in vitro* substrates including organoids and cell lines [23]. Use of structure-based approaches dates back the discovery of X-ray crystallography and nuclear magnetic resonance (NMR) spectroscopy; tools which determine the two- or three-dimensional structure of small molecules and their biological targets. These structure-based technologies have been a cornerstone of drug repurposing in medicinal chemistry [24]. In contrast, transcriptomic signature-based drug discovery is a more recent approach that classifies drugs based on their transcriptomic signatures [25]. Both approaches largely rely on public databases of drug structure and transcriptional signatures. These data may inform drug repurposing by incorporating information related to precise structure/signature-and-function relationships. In this section we discuss these two *in silico* methods with specific emphasis on the signature-based approach and the development of data science methods for its advancement. A summary of these different approaches and their respective tools is summarized in Table 3.

**Table 3.** Glossary of Databases and Software Applications

| **Application** | **Brief Definition** | **Website URL, Notes [Reference]** |
| --- | --- | --- |
| **AutoDock** | Automated docking software used to predict how candidate drugs bind to receptors. | http://autodock.scripps.edu. |
| **BANDIT** | Drug interaction targets software used to determine what receptor or enzyme interacts with prospective drugs. | Algorithm [99] |
| **CauseNet** | A platform utilized for drug repositioning based on causal network between drugs and disease. | Algorithm [73] |
| **CMap** | Reference database of gene expression profiles in cell lines exposed to chemical perturbagens. | https://www.broadinstitute.org/connectivity-map-cmap. |
| **DDBJ** | Nucleotide sequencing database. It is the only databank that is certified in Asia. | https://www.ddbj.nig.ac.jp/index-e.html. |
| **DOCK** | Virtual screen software used to predict and simulate the binding of small molecules with receptors. | http://dock.compbio.ucsf.edu. |
| **EBI** | Nucleotide sequencing database. Consider as the primary source for bioinformatic data in Europe. | https://www.ebi.ac.uk. |
| **GEO** | An international public repository database. Utilized to query, download experiments, and curated gene expression profiles. | https://www.ncbi.nlm.nih.gov/geo/. |
| **Glide** | Docking software that provides highly accurate orientation and positional space of the docked ligand upon adding chemical perturbagens. | Algorithm [40] |
| **LINCS** | A data portal that provides a repository of signatures from diverse cell types against variety of perturbing agents. | http://www.lincsproject.org. |
| **Merck Index** | Encyclopedia of a group of chemical substances and biological molecules grouped and published online by the Royal Society of Chemistry. | https://www.rsc.org/merck-index. |
| **ModBase** | A comparative protein structure models database. | https://modbase.compbio.ucsf.edu/modbase-cgi/index.cgi. |
| **National Cancer Institute's Drug Dictionary** | A library of technical definitions and synonyms of drugs used to treat cancer or cancer related conditions. | https://www.cancer.gov/publications/dictionaries/cancer-drug |
| **NCBI** | A house of distinct databases relevant to biotechnology and biomedicine | https://www.ncbi.nlm.nih.gov |
| **PharmGKB** | A database responsible for aggregation, curation, integration and dissemination of data pertaining to human genetic variation to drug response. | https://www.pharmgkb.org |
| **PDB** | A database for large biological molecules that are presented in a three-dimensional structure. | https://www.rcsb.org |
| **PubChem** | A database of chemical molecules and there activates against biological assays. | https://pubchem.ncbi.nlm.nih.gov |
| **RosettaDock** | A database for protien-protien docking interaction. | https://www.rosettacommons.org |

Abbreviations: BANDIT, Bayesian ANalysis to determine Drug Interaction Targets; CMap, Connectivity-Map Database; DDBJ, DNA Data Bank of Japan; EBI, European Bioinformatics Institute; GEO, Gene Expression Omnibus; LINCS, Integrated Network-based Cellular Signature; NCBI, National Center For Biotechnology Information; PDB, Protein Data Bank; ZINC, undefined name of non-commercial drug database.

### Structure based approaches

Structure-based approaches for repurposing rely on the principle of shape complementarity between the target protein and candidate molecule, and thus require the 3D structure of the target protein be resolved either by X-ray crystallography, NMR spectroscopy, or computational techniques, such as homology modeling [24]. With rapid advances in structural biology, 3D structures for most druggable proteins are available from the Protein Data Bank [26] or databases of pre-computed models, such as ModBase [27]. Computational docking and virtual screening approaches can then estimate the shape complementarity and binding affinity between the target and a ligand [28]. A typical docking simulation is performed for a large number of candidate compounds in order to rank them according to their predicted binding affinity and select top hits for further assessment and validation. This approach can accelerate drug repurposing by facilitating rapid identification of lead candidates by using a library of drug molecules, such as national cancer institutes (NCI) drug dictionary [29], library of integrated network-based cellular signature (LINCS) [30], ZINC (a non-commercial database of commercially available compounds for virtual screening) [31] or previously approved drugs. To speed up the computation, a docking program requires a computational cluster, or a distributed computing platform and virtual screening pipelines integrated with chemoinformatic analyses.

Conceptually, docking simulations involve two main components: sampling and scoring [32,33]. Sampling algorithms are used to find plausible conformations of the receptor-ligand complex, while scoring functions are required to estimate relative binding affinities and rank ligand poses (conformations of the receptor-ligand complex) [34]. This search through the space of possible conformations of the receptor-ligand complex can be computationally expensive and involves using various optimization techniques, such as a Monte Carlo simulation, simulated annealing, or genetic algorithms [35]. In order to provide the basis for scoring and ranking, atomic force fields and simplified solvation potentials are typically combined into empirical scoring functions that introduce many approximations to describe both intra- and inter-molecular interactions in the system, as well as to estimate the strength of interactions between the ligand and receptor [36]. As a result, different scoring functions may introduce distinct biases that have to be taken into account when selecting one of the available docking methods [37].

Stimulated by both methodological advances and fast changes in computing architectures, docking methods have improved considerably over the last decade. There are many software packages now available for computational docking (of small molecules) that may be used for docking simulations and drug repurposing, including AutoDock [38], DOCK [39], Glide [40], and RosettaDock [41]. Benchmarking of docking packages suggests that no single method consistently outperforms other approaches [42]. Therefore, different targets may require different combinations of methods, potentially enhanced by re-scoring approaches, including those using transcriptional signature-based approaches, to further limit significant false positive and false negative rates observed in docking studies [43]. Considering a consensus approach utilizing multiple programs offers a viable strategy to more reliably identify candidate drugs that are true binders of specific targets [42].

### Signature based approaches

Disease or drug-mediated alterations in mRNA expression can be used to define unique molecular signatures (Table 1) [23]. A compilation of molecular signatures allows for the selection of drugs for a particular disease signature by way of the signature reversion principal, which assumes that if a drug-induced transcriptional signature is similar or dissimilar to a disease signature, then that drug may restore or reverse the disease phenotype, respectively [44].

Disease-associated transcriptomic data are either self-generated in laboratories or are readily available from public repositories such as the National Center for Biotechnology Information (NCBI), European Bioinformatics Institute (EBI), or the DNA Data Bank of Japan (DDBJ). However, the data for drug-induced transcriptomic responses is not readily available and is complicated by several factors. On the clinical side, there is a paucity of systematic studies testing the effect of drugs and doses in the human subjects. In addition, there is a lack of drug treatment associated demographics in the available omics datasets. Further, on the pre-clinical and experimental side, there is a huge cost involved in generating transcriptomic profiles of millions of molecules, and the lack of standardized animal models and doses in which the drugs are tested prevents meaningful analyses. Circumventing these issues requires a systematic and cost-effective approach and repository to generate, assemble, and analyze such parameters across drugs.

The Broad Institute first piloted a gene expression profile compendium of pharmacological perturbagens, leading to the generation of the connectivity-map (CMap) database (Table 1) [45]. Initially, CMap was designed to utilize Affymetrix GeneChip microarrays to generate transcriptome-wide signatures by testing 164 small molecules on four cancer cell-lines (MCF7, PC3, HL60, and SKMEL5) [25]. To scale up the workflow and more deeply characterize a given perturbagen’s cellular signature, a new assay, the L1000, was developed [46]. The assay utilizes an optically addressed microsphere and a flow-cytometric detection system to measure gene expression levels of 978 landmark genes (nearly 1000, thus the name L1000) along with 80 control genes with invariable expression across different experiments [46]. The landmark genes provide a reduced representation of the full transcriptome derived from 12,031 gene expression profile available at GEO. The L1000 landmark genes were used to infer the expression of 11,350 other genes of the remaining transcriptome [47]. The inferred genes had high degree of similarity with the profile generated from RNAseq. The assay dropped the cost of generating the transcriptomic profile to approximately two dollars per sample and propelled the expansion of signature generation [47]. Now called the Library of Integrated Network-Based Cellular Signatures (LINCS), there are over one million L1000 profiles for more than 19,000 small molecules in >100 cell lines. Each small molecule is profiled in triplicate following treatment for 6 through 24 hours [47]. A number of FDA approved small molecules (~2500/19000) with known mechanism of action, protein targets, pharmacological classification, and clinical indication were profiled in 9 cancer cell-lines (Table 4). Data from these molecules, called “Touchstone,” are used as a reference to functionally annotate users query signatures by correlating implicated genes and pathways. The remaining uncharacterized small molecules profiled on >100 cell lines are termed “Discovery” datasets and may be annotated by connecting them with the Touchstone dataset. More L1000 transcriptome data is actively being generated from six different LINCS centers [48], as well as phosphoproteins signatures from the same cell culture studies (called P100)[30].

**Table 4.** Primary cell lines in the LINCS database

| Name | Cell type | Number of signatures associated with: |  |  |  |
| --- | --- | --- | --- | --- | --- |
|  |  | CGS, KD | Drug perturbagen | Overexpression | Proteomic |
| A375 | Malignant lung melanoma | 3,721 | 1,823 | 1823 | 183 |
| A549 | Non-small cell lung carcinoma | 3,673 | 9,186 | 591 | 184 |
| HEPG2 | Hepatocellular Carcinoma | 3,341 | 4,622 | 999 | 0 |
| HCC515 | Non-small cell lung Adenocarcinoma | 2,404 | 5,382 | 899 | 0 |
| HT29 | Colorectal Adenocarcinoma | 3,354 | 10,193 | 661 | 0 |
| MCF7 | Breast Adenocarcinoma | 5,169 | 18,846 | 639 | 156 |
| PC3 | Prostate Adenocarcinoma | 5,288 | 13,532 | 818 | 90 |
| VCAP | Metastatic Prostate Cancer | 3,636 | 14,751 | 608 | 0 |
| YAPC | Pancreatic Carcinoma | 53 | 4,440 | 0 | 181 |

Source: https://lincs.org; CGS, Consensus Gene Signature; KD, Knockdown; LINCS, Library of Integrated Network Based Signatures. Cell lines included in table consist of core LINCS cell lines comprising the largest subset of signatures available.

The standard approach for utilizing LINCS resources involves using disease- or phenotype-associated differentially expressed genes as a query against the drug signatures. Connectivity (or similarity) scores are then computed based on nonparametric, rank-based, pattern-matching algorithms [49], which identify a disease-inducing or therapeutic drug based on drug-disease similarity or dissimilarity, respectively. This connectivity score reflects the extent to which a drug modifies a disease-associated gene, without necessarily considering the magnitude of that change. To mitigate this issue, connectivity scores are normalized to account for global differences in cell and perturbation type. In addition, to compare the observed normalized connectivity score to all others in the LINCS repository, a percentile score ranging from −100 to +100 (called tau) is calculated. Since the reference to calculate the percentile is fixed, the tau score can be used to compare the results across many queries. In such an instance, a connection with significant p-value and but low tau would suggest a promiscuous drug with non-unique connections, while those with high tau scores would suggest a drug more specific to the disease signature. Several other approaches [50,51] and tools [52,53] exist to evaluate drug-drug and drug-disease similarity comparisons. A well-conceived article by Zhou et al. provides a systematic evaluation of a variety of these approaches [54].

Finally, compared to structure, the transcriptomic signature of a drug is highly variable. This is an inherent issue related to the present method where drugs are treated in various cell types, usually with three or more biological replicates. To overcome this challenge, as a final step towards signature-based data generation, the LINCS consortium is now generating consensus signatures, which are consistent (and thus comparable) across different molecules.

## III. Informed repurposing using data integration

The success of any drug development or repurposing approaches is contingent upon its potential to characterize the phenotype of interest and elucidate a mechanism of action. Signature-based approaches, being associated with gene expression, can make full use of expanding omics-based technology, data analysis, and integration workflows. For instance, knowledge can be incorporated from genome wide association studies (GWAS), disease biomarkers, and biological pathways associated with a disease state to filter for precise and causal drug candidates. In addition to omics-based filtering, current drug repurposing approaches involve integration of both structure and signature-based approaches with advanced data mining and machine learning methods. In the following section we discuss data integration efforts for more informed drug repurposing.

### GWAS based gene signatures for repurposing

Single nucleotide polymorphisms (SNPs) are the most common cause of variation in the human genome [55]. SNPs are single base pair changes that occur mostly in non-coding regions of the genome and may have a biological or functional contribution towards disease states. SNPs may impart disease risk by changing the affinity of a transcription factor for DNA binding site, the stability of the transcript, or the amino acid sequence of the translated protein [56]. Neuropsychiatric disorders have diverse risk alleles, where genomic variations, including SNPs, confer susceptibility to developing the disorder [57]. GWAS compares SNP alleles between the cases and controls to characterize disease-associated variations explained by SNPs. These genomic variations provide information on the genetic and biological underpinnings of a disease and represent an additional approach to target causal genes and gene-products for drug repurposing.

Several studies have used GWAS-based results as a disease signature to repurpose drugs for an array of neuropsychiatric disorders (Table 5). For example, a meta-analysis of 796 GWAS studies filtered 991 genes identified by GWAS [58,59]. 21% (212/991) of these genes were considered targetable by small molecules and 47% (469/991) were considered biopharmable, that is genes annotated as having a peptide product or containing a transmembrane domain. The list of genes returned from their analysis was enriched compared to those derived from the entire genome, which contains 17% drug-target for small molecules and 38% biopharmable genes. This study provides a strong rational for a GWAS based drug-discovery and repurposing pipelines [58].

**Table 5.** GWAS-identified drugs which may be repurposed for each indicated disorder

| Drug | MOA | Original Indication |
| --- | --- | --- |
| **Drugs repurposed for Alzheimer's Disease [129–131]** |  |  |
| Naftidrofuryl# | Vasodilator; selective antagonism of 5-HT<sub>2</sub> receptors | Peripheral Artery Disease, Intermittent Claudication |
| Vinpocetine | Synthetic *vinca* alkaloid, posited to inhibit Na<sup>+</sup> channels, cyclic nucleotide phosphodiesterase, and IκB kinase | Dietary supplement for cognition |
| Celecoxib, Naproxen, Meclofenamic Acid | NSAIDs, inhibition of cyclooxygenase-1 and -2 | Anti-pyretic, anti-inflammatory, and analgesic |
| Gemtuzumab ozogamicin | Monoclonal antibody against CD33 | Acute Myeloid Leukemia |
| Omalizumab | Recombinant antibody against IgE, inhibition of mast cell degranulation | Allergic Asthma |
| Leflunomide | Pyrimidine synthesis inhibitor, inhibition of T-cell activation | Rheumatoid Arthritis |
| **Drugs repurposed for Major Depressive Disorder [129,132]** |  |  |
| Pregabalin, gabapentin, and nitrendipine | Voltage-gated calcium channel antagonist | Neuropathic pain, adjunct therapy for epilepsy; hypertension, angina |
| Alizapride; mesoridazine | Dopamine D<sub>2</sub> receptor antagonists | Anti-emetic; neuroleptic for Schizophrenia |
| Levonorgestrel, Diethylstilbestrol | Synthetic progesterone; non-steroidal estrogen receptor agonist | Contraception and hormone therapy |
| Papaverine | Phosphodiesterase inhibitor | Vasodilator |
| Scopolamine | Muscarinic antagonist | Anti-emetic, motion sickness |
| Ketoconazole | Inhibition of steroidogenesis, cortisol lowering agent | Anti-fungal, Cushing's syndrome |
| Arcaïne sulfate#, ifenprodil, cycloserine | Glutamatergic modulation: NMDA receptor antagonists; cycloserine is a partial agonist. | Arcaïne sulfate and Ifenprodil is implicated in basic science research; cycloserine is an antibiotic |
| Risperidone; Sulpiride | Serotonergic and dopaminergic antagonists; Dopamine D<sub>2</sub> receptor antagonist | Antipsychotic medication for Bipolar, Schizophrenia |
| **Drugs repurposed for Schizophrenia [132–135]** |  |  |
| Fluticasone | Glucocorticoid receptor agonist | COPD and Asthma |
| Resveratrol | Suppression of NF-Kappa-B activation in Herpes Simplex Virus infected cells | Antineoplastic, anti-inflammatory |
| Zonisamide | Sodium and T-type calcium channels blocker, weak inhibition of carbonic anhydrase or monoamine oxidase B, and allosteric modulator of GABA receptor | Antiepileptic, antiparkinsonian and adjunctive therapy for bipolar and anxiety disorders |
| Bevacizumab | It is a monoclonal antibody targeting vascular endothelial growth factor A | Antineoplastic |
| Verapamil | Calcium channel blocker | Hypertension, arrhythmias, angina |
| Cinnarizine | Calcium channel blocker | Antihistamine, antiemetic, and antagonist of dopamine D2 receptors |
| Varenicline | Nicotinic agonist | Smoking cessation |
| Galantamine | Allosteric modulator of nicotinic receptors and an acetylcholinesterase inhibitor | Smoking cessation; Alzheimer's disease |
| Dutogliptin, Alogliptin | Dipeptidyl peptidase 4 inhibitors | Type 2 Diabetes Mellitus |
| Atorvastatin | Competitive inhibitor of HMG-CoA reductase | Dyslipidemia |
| Metoclopramide; Trifluoperazine | Dopamine antagonist | Diabetic gastroparesis; typical antipsychotic |
| Neratinib | Tyrosine kinase inhibitor | Antineoplastic |
| Nifedipine | Calcium channel blocker | Hypertension |
| Xanomeline<sup>#</sup> | Muscarinic acetylcholine receptor and serotonergic agonist | Antipsychotic |
| Mecamylamine | Nicotinic antagonist | Hypertension |
| Memantine | NMDA receptors antagonist | Alzheimer's disease; neurological disorders |
| **Drugs repurposed for Epilepsy [136]** |  |  |
| Vinorelbine | Third generation *vinca* alkaloid, microtubule inhibitor | Antineoplastic |
| Nornicotine<sup>#</sup> | Acetylcholine receptor agonist | Insecticide, no indication. |
| Equol | An estrogen receptor agonist | Antineoplastic |

A criticism of GWAS is that it may identify SNPs with spurious (or “passenger”) associations to the disease or disorder within a region of the genome instead of pinpointing *bona fide* associations [60,61]. However, there are techniques available that permit identification of causal genes. Technologies such as Transethnic GWAS (studies prioritizing candidate genes across diverse populations) [62], copy number variant analysis (identifying variations in the number of gene copies) [63], quantitative trait loci analyses (mapping complex phenotypes to a chromosomal locus) [64], imputed gene expression profiles [65], and epigenomic methods improve identification of causal disease genes [66], narrowing down “druggable” targets [67]. These methods are typically deployed in conjunction with network- and pathway-informed approaches.

### Data driven drug repurposing

The network-based approaches to drug-discovery and repurposing largely builds upon the guilty-by-association principle, which assumes that closely related drug signatures in a network may share gene expression signatures and are likely to have related function [68,69]. Thus, efficacy of an unknown drug may be inferred based on its proximity to drugs with known function in a defined network. To date, this approach has only been widely applied to FDA approved drugs by examining drug-disease interaction networks. However, this approach has significant limitations, as it typically centers on a small number of interactions whose functional properties cannot be extrapolated to the rest of the network. In effect, it only encompasses outliers (representing a few strong connections) whose function is not necessarily generalizable [70]. To mitigate this limitation, signature-based pathway information [71] and data from other networks (including drug-drug, target-target, disease-disease) may be integrated to create a more heterogeneous network [72].

Pathways or ontologies functionally represent related genes and provide summative information for their interactions. Pathways have been extensively utilized to explore cross talk between different biological process, gene-gene interactions, molecular mechanisms underlying a disease, and disease causality. For example, a network-based approach was combined along with the disease-associated pathways [73]. This platform, called CauseNet, mimics a manual pathway analysis for drug repurposing and works via a multilayer network construct linking drug to target, target to pathway, pathways to gene, and gene to disease. The transition likelihood from one link to another is learned using statistical methods. A novel indication for a drug is predicted using a maximum likelihood estimation. In a cross validation test, the approach showed high performance (AUC=0.859) in predicting novel indications for known repurposed drugs [73].

Using a similar strategy, a heterogeneous network was created by combining information from disease-disease, drug-drug and target-target networks [74]. This approach outperformed similar methods utilizing only drug-drug similarity and permits prediction of drug-target and drug-disease relationship. This approach was recently extended using a Bi-Random walk algorithm to predict novel indications of an existing drug [75]. The algorithm improved the predicted drug-disease similarity in two parts. First, it adjusts the weak and uninformative drug-disease similarity by randomly permuted correlation analysis. Second, after adjustment, rather than correlation, any two drugs were considered similar based on number of shared (common) drugs between the two drugs. The approach performed better than network-based algorithms (AUC = 0.91) [75].

Ongoing work is also merging data from both structure and signature-based approaches to increase efficiency and selectivity. For example, a database of 4296 compounds was created using signature profiles derived from 60 human tumor cell lines [76]. The pairwise similarity of each signature was computed and correlations between compounds greater than or equal to 0.75 were considered robust. Next, the chemical similarity for these compounds was computed and those with similar structural and signature profiles were identified as possibly having the same molecular target, suggesting new indications for repurposed compounds. A related attempt to integrate signature and structure-based approaches for drug repurposing generated 62 vectors (per compound) for 147 drugs that are inhibitors of cruzipain, a parasitic cysteine protease [77]. The model trained using these vectors was then used to predict cruzipain inhibiting activity for 5000 compounds from Merck Index 12^th^ database [78]. Docking simulation of compounds predicted by the model was performed to assess their bioactivity against the cruzipain protein. Such integration of signature and structural approaches is a promising and robust advance for drug repurposing.

### Text mining to filter relevant information

The structural and transcriptomic signatures of a drug may suggest hundreds of compounds with similar effects; in such an instance, filtering or enriching for the most promising candidates can be challenging. However, combining critical information regarding a drug’s family, pharmacology, toxicity, protein target, and targeted pathways, as well as structural and signature-based similarity, may enhance the drug-repurposing workflow [79]. Text mining combines information from thousands of documents and articles to deliver new meaning and possible answers to complex questions [80]. Text mining has been used prolifically in the medical field and in conjunction with data-driven drug repurposing approaches [81]. A typical biological text mining effort involves four steps: 1) information retrieval, including parsing of relevant information from large data sources; 2) biological name entity recognition, with identification of valuable biological concepts using controlled vocabularies, and the last two steps; 3) biological information extraction; and finally, 4) biological knowledge discovery, which involves extracting useful biological information and constructing a knowledge graph, a compilation of interlinked descriptions of objects [82].

Text mining was recently utilized in combination with network analysis to explore disease-protein and drug-protein relationships [83]. Using known proteins implicated in AD, a protein-protein interaction network was curated. The list of proteins in this network was used as query to text mine drug related information from PubMed abstracts [83]. This query returned a list of 1,249 possible AD related drugs. Then, drug-target similarity scores were calculated to assess for biological relevance. Their approach led to the identification of diltiazem and quinidine, prescribed for hypertension and cardiac arrythmias, respectively, as possible drugs of interest in treating AD. This work highlights the potential of text mining as an approach to identify previously unknown patterns memes and to generate hypothesis from accumulating text sources and databases.

## IV. Beyond Traditional Drug Repurposing: Inclusion of Diverse Omics Datasets

Using signature-based drug repurposing approaches for a given disorder generates surprising and often novel insight into disease pathology. While acquiring new knowledge of disorders is crucial for medicine and the scientific community, such techniques also allow for an improved understand of the drugs used to treat them. As we grow our attempts to intelligently repurpose drugs, we find that we may discover new knowledge related to drug classifications, adverse drug reactions, mechanisms of action, as well as the genetic predispositions that drive drug sensitivity and resistance. These domains and others are being deployed and leveraged for integrative drug repurposing.

### Drug classification

The anatomical therapeutic chemical (ATC) system created by world health organization classifies drugs into different groups based on the organ system they target, as well as their chemical, pharmacological and therapeutic properties [84]. Such a classification serves as a tool to improve drug utilization and development. Transcriptomic signatures of drugs may also be used to develop drug classifications. A novel machine learning based drug classifier was recently developed, which focuses primarily on the drug characteristics and class extracted from ATC, instead of relying on drug-disease similarity [85]. The classifier involves calculating average similarity between drugs to predict drug class using three features: gene-expression signatures, chemical structure, and known common targets calculated based on human protein-protein interaction networks. Each feature by itself had minimal performance in the classifiers. However, the performance increased to an accuracy of 78% when all three features were used in combination to predict the ATC class of 281 drugs. This method may be even more powerful than these high accuracy levels suggest. Subsequent analyses suggested that these “misclassified” drugs (i.e., the remaining 22% of these drugs) are more accurately conceptualized as being “reclassified.” In other words, this particular classifier goes beyond simple validation of the ATC classification system and has an ability to suggest new drug classifications that may prove more useful than the original drug classification paradigm. The classifier could predict most known drugs into new therapeutic classes, which were consistent with several literature reports. In a similar study, chemical-chemical similarity and interaction were used as features to predict the ATC class of 3,833 drugs. This approach predicted drug class with 73% accuracy, substantially higher than 7% accuracy using prediction by chance [86].

The ATC also has important limitations. It has incomplete drug coverage, and the classification rubric presently involves 14 levels, with each level further divided into subgroups; this occasionally leads to singleton drug classes with only one drug [87]. In addition, the classification of a new drug using ATC is a tedious process, involving a formal request for classification from researchers to world health organization [88]. Data driven drug-classification using drug signatures will improve the utility and accuracy of the ATC and related databases.

### Adverse Drug effects

An adverse drug reaction (ADR) can be defined as a harmful reaction to a drug, causing injury with a clear causative link to the drug being administered [89–91]. ADRs are a major concern for both drug development and public health, and failure to identify ADRs can lead to significant morbidity and economic loss. However, ADRs may also help in understanding drug-disease phenotype connections. Typically, predicting an ADR was considered a binary classification problem where the chemical and biological aspects of the drugs were used to predict the presence or absence of adverse effects. Taking advantage of this possible connection, SEP-L1000, a machine learning based classifier, was developed to predict ADRs of over 20,000 small molecules available in the LINCS database [92]. The steps in developing the classifier involved gathering drug associated data (features) from multiple resources including, but not limited to, structure from PubChem [93], signatures from LINCS, as well as side effect data from Drug Side Effect Resource [94] and PharmGKB (a resource for ADRs of FDA approved drugs) [95]. To prioritize the most predictive features for each ADR class, feature selection was performed using a regularized logistic regression model. After selecting the top 50 predictive features, classification algorithms were applied to train each ADR class [96]. Importantly, benchmark metrics showed that gene expression signature was the best predictive feature and was used further to associate each drug’s ADR with the gene ontology-based pathway networks. This novel approach performs better than target-based binary classification for predicting ADRs and is scalable. Further, incorporation of gene ontology in the model may provide new insights about a drugs mechanism for ADRs.

### Drug Mode of action

Mode of action (MOA) refers to specific drug-target interaction through which the pharmacological effect of drug is observed [97]. Beside the traditional structure based molecular docking approaches of identifying the drug-target interaction, newer and faster computational modeling approaches leveraging a drug’s transcriptome signatures are being developed [98]. For example, a Bayesian machine learning approach (called BANDIT: Bayesian ANalysis to determine Drug Interaction Targets) was developed to identify drug targets. BANDIT combines LINCS resources, as well as data for drug structure, growth inhibition, side effects, known targets, and bioassay results. With integration of these predictors, BANDIT provides an improvement in predicting drug targets over other similar approaches [99]. The method calculates a pairwise similarity score for each predictor for all drugs with both known and unknown shared targets. To assess the degree of each data type’s ability to separate the pairs groups, a Kolmogorov–Smirnov test was used. Their results indicated that structural similarity; bioassay and growth inhibition assays had the strongest differentiation statistic, while the transcriptional and adverse effects had the weakest. Benchmarking showed that BANDIT had overall accuracy of ~90% in predicting the mode of action of a drug. Determining MOA *in silico* allows for additional characterization, rather than purely relying on biochemical bench-work, with high accuracy that may increase the drug-development pipeline.

### Drug resistance

Drug resistance, a decrease in effectiveness of a therapeutic to treat a disease condition, is one of the primary obstacles facing drug design and discovery today. In cancer, somatic mutations have considerable impact on drug resistance [100]. To address this concern, expression-based variant-impact phenotyping was developed, an approach which compares wild-type and mutant alleles of the same gene’s transcriptional signatures to deduce functional role(s) of that specific mutations [101]. This approach also segregates the impact of somatic mutation on drug resistance into consequential mutations, called drivers, and non-consequential mutations, called passengers, and allows omics scale determination of the contribution of somatic mutations to cellular functions. It follows that for a successful drug repurposing, gene expression profiling of a drug should distinguish the contribution of somatic mutations to perturbagen signatures. As the number of molecule-specific transcriptional signatures in LINCS grows providing signatures associated with gene variants, the ability to develop drugs that consider and overcome drug resistance will improve. Recent work highlights this approach.

LINCS resources and transcriptional profiles of biopsies from patients before, during and after relapse were used to assess the variable efficacy of chemotherapies, including MEK and BRAF inhibitors which have been suboptimal in treating tumors [47]. A comparison of mutations in the tumors with profiling of existing perturbagen, knockdown, and overexpression signatures in LINCS cancer cell lines showed a strong negative correlation with MAP kinase signaling, suggesting a re-activation of the MAPK pathway in these patients during relapse [47]. Although the contribution of somatic mutations to psychiatric disorders is not well characterized, this method could be leveraged to study genomic variants associated with treatment resistant depression, schizophrenia, or other psychiatric disorders, highlighting an opportunity to develop more effective pharmacotherapies.

### Drug permeability

Influenced by lipophilicity, size, charge, molecular weight, and hydrogen bonding capacity, permeability of drugs across membranes is key parameter influencing its absorption, distribution, and elimination across blood brain barrier [102]. Based on the available 2D or 3D structure of the drug, permeability across membranes can be estimated [103]. One approach to this challenge utilizes the LINCS L1000 landmark genes. Based on the principal that a more permeable compound will tend to induce larger changes in transcript expression, the expression levels of the L1000 genes can be used as a proxy of cellular permeability for a compound or drug in the LINCS database. Transcriptional activity score (TAS), a proxy for molecules cellular permeability and activity, is estimated for each compound in the LINCS database. TAS ranges between 0 and 1, and is a geometric mean of signature strength (number of differentially expressed landmark gene with absolute z-score >2 for a given compound) and replicate correlation (correlation between biological replicate of a compounds L1000 profile) normalized by number of L1000 genes. So far the use of TAS is limited to drug repurposing efforts against antimicrobial agents [47,104] and its potential to predict complex blood brain barrier is uncertain. However, since TAS is associated with gene signatures it may be linked to ontology and used as a predictor in complex models to determine drug activity.

### Generation of seed gene knockdown signatures

Studies investigating the pathophysiology of CNS disorder using postmortem brain or model systems often focus on one or a few candidate genes. Such hypothesis- or candidate-driven research often fails to account for the complexity of CNS diseases, as well as the heterogeneity of biological processes. However, using so-called “seed genes” chosen based on findings related to specific candidate genes or pathways can be an entry point for bioinformatics analyses, connecting candidate-based studies with drug repurposing. Combining the changes in expression of a small number of genes in a disease state along with LINCS gene knockdown and/or overexpression signatures permits interrogation of the LINCS perturbagen database for compounds that reverse or simulate consensus “seed gene” signatures. This approach is built on the assumption that seed gene-specific signatures would reflect disease-associated changes at the network level [105].

A recent study highlights the potential of the seed gene approach. Knockdown signatures of glycolytic genes decreased in pyramidal neurons in schizophrenia were used to generate clusters of genes to probe the LINCS database [15,106,107]. Using LINCS, drugs and compounds were identified that reversed the consensus seed gene clusters for the implicated glycolytic genes. Pioglitazone, a synthetic ligand for Peroxisome proliferator-activated receptor gamma, a nuclear receptor and a member of the thiolazinedione drug family, showed significant reversal of these bioenergetic profiles. Pioglitazone is a well-characterized regulator of bioenergetic function, including lipid homeostasis, adipocyte differentiation, and insulin sensitivity. It also increases expression of glucose transporters. As a confirmation study, decreased levels of the glucose transporters GLUT1 and GLUT3 were found in schizophrenia as well as in the Grin1KD mouse, an animal model of developmental disorders [105]. Treatment of Grin1KD mice with pioglitazone for 1 week improved executive function. Start-to-finish this work illustrates the potential for repurposing FDA approved drugs, starting with candidate or seed genes, followed by bioinformatics-driven identification of drug candidates, and finally testing in animal models [105].

### V. Caveats

Compared to drug repurposing efforts realized by empirical methods, there are a few limitations associated with signature-based repurposing, which may limit its clinical relevance. First, most studies are still in the exploratory stage and results are limited to FDA approved drugs with known mechanisms of action. Expanding these methods to include yet unclassified and untested small molecules from compound libraries is ongoing, but validating biological significance remains incomplete. This may be due to lack of animal models for the disease in question or poor simulation of disease in the animal model. Novel *in-vitro* organ-on-chip [108] or organoid [109] based drug screening techniques may be integrated with the present *in silico* approaches to circumvent some of these limitations. Second, the clinical and epidemiological techniques supporting empirical drug repurposing strategies have revealed important information related to dose compatibility and disease comorbidity. These techniques have not been sufficiently incorporated into predictors for machine learning models used for signature-based drug repurposing. Integration of these data can potentially increase the discovery of new indications ultimately improving drug-repurposing efforts.

### VI. A Path forward

Historically, many causal theories of psychiatric disorders were based on or informed by unintended discoveries of medications with efficacy for a specific disorder. For instance, the use of iproniazid and imipramine contributed to the monoamine hypothesis of depression [110]; fluoxetine led to the discovery of serotonin in relation to depression [111]; ketamine, an anesthetic, contributed to the glutamate hypothesis of depression [112] and lithium, a receptor-naive salt, highlighted the role of neuronal cell homeostasis and intracellular signaling in bipolar disorder [113]. Currently, bioinformatics tools offer windows of opportunity to explore new mechanisms of psychiatric disorders.

A recurrent theme that emerges in the aforementioned approaches of signature-based repurposing and knowledge discovery is integration, both for datasets and bioinformatic tools. A feature (or predictor) defined by a particular approach by itself has limited power to predict the outcome (new indication or knowledge). Combining an ensemble of approaches along with computational power and statistical innovation has shown promising results. In this regard, several future avenues should be considered. First, with more recent expansion of transcriptomic technologies to single cellular resolution, data-centric repurposing approaches can now integrate single-cell-transcriptomic data to refine mechanism pathophysiology to a cellular level. Second, lead drugs from signature-based repurposing may be used as skeletons to design or optimize new molecules using chemoinformatics approaches [114]. Third, transcriptomic signatures of drugs can be associated with gene ontology, enabling pathway-specific drug repurposing. Taken together, integration of datasets and bioinformatics tools and platforms offers a promising avenue for advancing the field of drug discovery via repurposing for psychiatric disorders.

### Drug Repurposing and Precision Medicine

Pharmacogenomics, a study of genomic influence in an individual’s response to drug, highlights inter-individual differences in drug response and contributes to developing precision medicine; a customized treatment decision tailored for an individual based on his/her genetic profile [115]. In psychiatry, the current practice in line with this idea includes repurposing drugs via off-label prescribing [116]. Possibilities exist to integrate the aforementioned-drug repurposing resources and approaches along with an individual’s genomic profile for a “precise-repurposing.” Thus far the major push in biomedical research at large for precise repurposing is from cancer genomics, where reference genomic profiles from cancer cell lines, solid tumors, leukemias, and stem cells are available and can be sequenced with relative ease [117]. In contrast, in psychiatry it is difficult to get brain tissues for obvious reasons, limiting access to the source of the pathophysiology for non-malignant, CNS disorders. However, inducible pluripotent stem cells (iPSCs), organoids, and other primary tissue sources are providing important new substrates that may be exploited for bioinformatic analyses. For example, iPSCs from a schizophrenia subject with the DISC mutation were used for an informatics-based study of kinase activity, identifying drug candidates that may reverse the cellular signatures found in these cells [118]. This study highlights the promise of personalized medicine, suggesting that integration of bioinformatics platforms, precision medicine, and drug repurposing may herald a paradigm shift for the field

## VII. Conclusions

In this review, we present the current status in the field for drug repurposing as it applies to CNS disorders, and more specifically for psychiatric illnesses. Transcriptional profiling represents an important recent advance for drug repurposing, and integration of traditional approaches with this promising technique show promise for advancing the field. Finally, there is considerable promise for deployment of personalized drug repurposing for psychiatric disorders, offering new avenues for translational research connecting big data analytics with the afflicted.

## Acknowledgements

This work was supported by NIMH MH107487 and MH121102.

## Reference

1. Pankevich DE, Altevogt BM, Dunlop J, Gage FH, Hyman SE. Improving and accelerating drug development for nervous system disorders. Neuron. 2014.

2. US Food and Drug Administration. New Molecular Entity (NME) Drug and New Biologic Approvals. 2014.

3. Lee HM, Kim Y. Drug Repurposing Is a New Opportunity for Developing Drugs against Neuropsychiatric Disorders. Schizophr Res Treatment. 2016.

4. Fava M. The promise and challenges of drug repurposing in psychiatry. World Psychiatry. 2018;17:28–29.

5. Hemphill CS, Sampat BN. Evergreening, patent challenges, and effective market life in pharmaceuticals. J Health Econ. 2012. 2012. https://doi.org/10.1016/j.jhealeco.2012.01.004.

6. Caban A, Pisarczyk K, Kopacz K, Kapuśniak A, Toumi M, Rémuzat C, et al. Filling the gap in CNS drug development: evaluation of the role of drug repurposing. J Mark Access Heal Policy. 2017. 2017. https://doi.org/10.1080/20016689.2017.1299833.

7. Leo RJ, Narendran R. Anticonvulsant Use in the Treatment of Bipolar Disorder: A Primer for Primary Care Physicians. Prim Care Companion J Clin Psychiatry. 1999. 1999. https://doi.org/10.4088/pcc.v01n0304.

8. López-Muñoz F, Shen WW, D’ocon P, Romero A, Álamo C. A history of the pharmacological treatment of bipolar disorder. Int J Mol Sci. 2018.

9. Bowden C. The effectiveness of divalproate in all forms of mania and the broader bipolar spectrum: Many questions, few answers. J Affect Disord. 2004. 2004. https://doi.org/10.1016/j.jad.2004.01.003.

10. Romoli M, Mazzocchetti P, D’Alonzo R, Siliquini S, Rinaldi VE, Verrotti A, et al. Valproic Acid and Epilepsy: From Molecular Mechanisms to Clinical Evidences. Curr Neuropharmacol. 2018. 2018. https://doi.org/10.2174/1570159×17666181227165722.

11. Serafini G, Howland R, Rovedi F, Girardi P, Amore M. The Role of Ketamine in Treatment-Resistant Depression: A Systematic Review. Curr Neuropharmacol. 2014. 2014. https://doi.org/10.2174/1570159×12666140619204251.

12. Fava M, Rush AJ, Wisniewski SR, Nierenberg AA, Alpert JE, McGrath PJ, et al. A comparison of mirtazapine and nortriptyline following two consecutive failed medication treatments for depressed outpatients: A STAR*D report. Am J Psychiatry. 2006. 2006. https://doi.org/10.1176/ajp.2006.163.7.1161.

13. Petersen T, Gordon JA, Kant A, Fava M, Rosenbaum JF, Nierenberg AA. Treatment resistant depression and Axis I co-morbidity. Psychol Med. 2001. 2001. https://doi.org/10.1017/S0033291701004305.

14. Serafini G, Pompili M, Innamorati M, Dwivedi Y, Brahmachari G, Girardi P. Pharmacological Properties of Glutamatergic Drugs Targeting NMDA Receptors and their Application in Major Depression. Curr Pharm Des. 2013. 2013. https://doi.org/10.2174/13816128113199990293.

15. Sullivan CR, Koene RH, Hasselfeld K, O’Donovan SM, Ramsey A, McCullumsmith RE. Neuron-specific deficits of bioenergetic processes in the dorsolateral prefrontal cortex in schizophrenia. Mol Psychiatry. 2019. 1 March 2019. https://doi.org/10.1038/s41380-018-0035-3.

16. Powell TR, Murphy T, Lee SH, Price J, Thuret S, Breen G. Transcriptomic profiling of human hippocampal progenitor cells treated with antidepressants and its application in drug repositioning. J Psychopharmacol. 2017. 2017. https://doi.org/10.1177/0269881117691467.

17. Williams G, Gatt A, Clarke E, Corcoran J, Doherty P, Chambers D, et al. Drug repurposing for Alzheimer’s disease based on transcriptional profiling of human iPSC-derived cortical neurons. Transl Psychiatry. 2019. 2019. https://doi.org/10.1038/s41398-019-0555-x.

18. Boldrini M, Underwood MD, Hen R, Rosoklija GB, Dwork AJ, John Mann J, et al. Antidepressants increase neural progenitor cells in the human hippocampus. Neuropsychopharmacology. 2009. 2009. https://doi.org/10.1038/npp.2009.75.

19. Malberg JE, Eisch AJ, Nestler EJ, Duman RS. Chronic antidepressant treatment increases neurogenesis in adult rat hippocampus. J Neurosci. 2000. 2000. https://doi.org/10.1523/jneurosci.20-24-09104.2000.

20. Moreira PI, Carvalho C, Zhu X, Smith MA, Perry G. Mitochondrial dysfunction is a trigger of Alzheimer’s disease pathophysiology. Biochim Biophys Acta - Mol Basis Dis. 2010.

21. Swerdlow RH, Khan SM. A ‘mitochondrial cascade hypothesis’ for sporadic Alzheimer’s disease. Med Hypotheses. 2004. 2004. https://doi.org/10.1016/j.mehy.2003.12.045.

22. Lionta E, Spyrou G, Vassilatis D, Cournia Z. Structure-Based Virtual Screening for Drug Discovery: Principles, Applications and Recent Advances. Curr Top Med Chem. 2014. 2014. https://doi.org/10.2174/1568026614666140929124445.

23. Iorio F, Rittman T, Ge H, Menden M, Saez-Rodriguez J. Transcriptional data: A new gateway to drug repositioning? Drug Discov Today. 2013.

24. Batool M, Ahmad B, Choi S. A structure-based drug discovery paradigm. Int J Mol Sci. 2019.

25. Lamb J. The Connectivity Map: A new tool for biomedical research. Nat Rev Cancer. 2007.

26. Berman H, Henrick K, Nakamura H. Announcing the worldwide Protein Data Bank. Nat Struct Biol. 2003.

27. Pieper U. MODBASE: a database of annotated comparative protein structure models and associated resources. Nucleic Acids Res. 2006. 2006. https://doi.org/10.1093/nar/gkj059.

28. Kitchen DB, Decornez H, Furr JR, Bajorath J. Docking and scoring in virtual screening for drug discovery: Methods and applications. Nat Rev Drug Discov. 2004.

29. Holbeck SL. Update on NCI in vitro drug screen utilities. Eur J Cancer. 2004. 2004. https://doi.org/10.1016/j.ejca.2003.11.022.

30. Koleti A, Terryn R, Stathias V, Chung C, Cooper DJ, Turner JP, et al. Data Portal for the Library of Integrated Network-based Cellular Signatures (LINCS) program: Integrated access to diverse large-scale cellular perturbation response data. Nucleic Acids Res. 2018. 2018. https://doi.org/10.1093/nar/gkx1063.

31. Irwin JJ, Shoichet BK. ZINC - A free database of commercially available compounds for virtual screening. J Chem Inf Model. 2005. 2005. https://doi.org/10.1021/ci049714+.

32. Huang SY, Zou X. Advances and challenges in Protein-ligand docking. Int J Mol Sci. 2010.

33. Ripphausen P, Nisius B, Bajorath J. State-of-the-art in ligand-based virtual screening. Drug Discov Today. 2011.

34. Morris GM, Lim-Wilby M. Molecular docking. Methods Mol Biol. 2008. 2008. https://doi.org/10.1007/978-1-59745-177-2_19.

35. Kellenberger E, Rodrigo J, Muller P, Rognan D. Comparative evaluation of eight docking tools for docking and virtual screening accuracy. Proteins Struct Funct Genet. 2004. 2004. https://doi.org/10.1002/prot.20149.

36. Rajamani R, Good AC. Ranking poses in structure-based lead discovery and optimization: Current trends in scoring function development. Curr Opin Drug Discov Dev. 2007.

37. Warren GL, Andrews CW, Capelli AM, Clarke B, LaLonde J, Lambert MH, et al. A critical assessment of docking programs and scoring functions. J Med Chem. 2006. 2006. https://doi.org/10.1021/jm050362n.

38. Goodsell DS, Morris GM, Olson AJ. Automated docking of flexible ligands: Applications of AutoDock. J Mol Recognit. 1996. 1996. https://doi.org/10.1002/(SICI)1099-1352(199601)9:1<1::AID-JMR241>3.0.CO;2-6.

39. Lang PT, Brozell SR, Mukherjee S, Pettersen EF, Meng EC, Thomas V, et al. DOCK 6: Combining techniques to model RNA-small molecule complexes. RNA. 2009. 2009. https://doi.org/10.1261/rna.1563609.

40. Friesner RA, Banks JL, Murphy RB, Halgren TA, Klicic JJ, Mainz DT, et al. Glide: A New Approach for Rapid, Accurate Docking and Scoring. 1. Method and Assessment of Docking Accuracy. J Med Chem. 2004. 2004. https://doi.org/10.1021/jm0306430.

41. Davis IW, Baker D. RosettaLigand Docking with Full Ligand and Receptor Flexibility. J Mol Biol. 2009. 2009. https://doi.org/10.1016/j.jmb.2008.11.010.

42. Yang JM, Chen YF, Shen TW, Kristal BS, Hsu DF. Consensus scoring criteria for improving enrichment in virtual screening. J Chem Inf Model. 2005. 2005. https://doi.org/10.1021/ci050034w.

43. Kim R, Skolnick J. Assessment of programs for ligand binding affinity prediction. J Comput Chem. 2008. 2008. https://doi.org/10.1002/jcc.20893.

44. Tan F, Yang R, Xu X, Chen X, Wang Y, Ma H, et al. Drug repositioning by applying ‘expression profiles’ generated by integrating chemical structure similarity and gene semantic similarity. Mol Biosyst. 2014. 2014. https://doi.org/10.1039/c3mb70554d.

45. Lamb J, Crawford ED, Peck D, Modell JW, Blat IC, Wrobel MJ, et al. The Connectivity Map: using gene-expression signatures to connect small molecules, genes, and disease. Science. 2006;313:1929–1935.

46. Peck D, Crawford ED, Ross KN, Stegmaier K, Golub TR, Lamb J. A method for high-throughput gene expression signature analysis. Genome Biol. 2006. 2006. https://doi.org/10.1186/gb-2006-7-7-r61.

47. Subramanian A, Narayan R, Corsello SM, Peck DD, Natoli TE, Lu X, et al. A Next Generation Connectivity Map: L1000 Platform and the First 1,000,000 Profiles. Cell. 2017. 2017. https://doi.org/10.1016/j.cell.2017.10.049.

48. Keenan AB, Jenkins SL, Jagodnik KM, Koplev S, He E, Torre D, et al. The Library of Integrated Network-Based Cellular Signatures NIH Program: System-Level Cataloging of Human Cells Response to Perturbations. Cell Syst. 2018.

49. Subramanian A, Tamayo P, Mootha VK, Mukherjee S, Ebert BL, Gillette MA, et al. Gene set enrichment analysis: A knowledge-based approach for interpreting genome-wide expression profiles. Proc Natl Acad Sci U S A. 2005;102:15545–15550.

50. Zhang SD. A simple and robust method for connecting small-molecule drugs using gene-expression signatures. BMC Bioinformatics. 2008. 2008. https://doi.org/10.1186/1471-2105-9-258.

51. Cheng J, Yang L. Comparing gene expression similarity metrics for connectivity map. Proc. - 2013 IEEE Int. Conf. Bioinforma. Biomed. IEEE BIBM 2013, 2013.

52. Zhang SD, Gant TW. sscMap: An extensible Java application for connecting small-molecule drugs using gene-expression signatures. BMC Bioinformatics. 2009. 2009. https://doi.org/10.1186/1471-2105-10-236.

53. Lee BKB, Tiong KH, Chang JK, Liew CS, Abdul Rahman ZA, Tan AC, et al. DeSigN: Connecting gene expression with therapeutics for drug repurposing and development. BMC Genomics. 2017. 2017. https://doi.org/10.1186/s12864-016-3260-7.

54. Zhou X, Wang M, Katsyv I, Irie H, Zhang B. EMUDRA: Ensemble of multiple drug repositioning approaches to improve prediction accuracy. Bioinformatics. 2018. 2018. https://doi.org/10.1093/bioinformatics/bty325.

55. Brookes AJ. The essence of SNPs. Gene. 1999.

56. Griffith OL, Montgomery SB, Bernier B, Chu B, Kasaian K, Aerts S, et al. ORegAnno: An open-access community-driven resource for regulatory annotation. Nucleic Acids Res. 2008. 2008. https://doi.org/10.1093/nar/gkm967.

57. Reich DE, Lander ES. On the allelic spectrum of human disease. Trends Genet. 2001.

58. Sanseau P, Agarwal P, Barnes MR, Pastinen T, Richards JB, Cardon LR, et al. Use of genome-wide association studies for drug repositioning. Nat Biotechnol. 2012.

59. Sanseau P, Agarwal P, Barnes MR, Pastinen T, Richards JB, Cardon LR, et al. Reply to Rational drug repositioning by medical genetics. Nat Biotechnol. 2013.

60. McClellan J, King MC. Genetic heterogeneity in human disease. Cell. 2010.

61. Boyle EA, Li YI, Pritchard JK. An Expanded View of Complex Traits: From Polygenic to Omnigenic. Cell. 2017.

62. Li YR, Keating BJ. Trans-ethnic genome-wide association studies: advantages and challenges of mapping in diverse populations. Genome Med. 2014;6:91.

63. McCarroll SA. Extending genome-wide association studies to copy-number variation. Hum Mol Genet. 2008. 2008. https://doi.org/10.1093/hmg/ddn282.

64. Damerval C, Maurice A, Josse JM, De Vienne D. Quantitative trait loci underlying gene product variation: A novel perspective for analyzing regulation of genome expression. Genetics. 1994. 1994.

65. Gusev A, Ko A, Shi H, Bhatia G, Chung W, Penninx BWJH, et al. Integrative approaches for large-scale transcriptome-wide association studies. Nat Genet. 2016. 2016. https://doi.org/10.1038/ng.3506.

66. Tak YG, Farnham PJ. Making sense of GWAS: Using epigenomics and genome engineering to understand the functional relevance of SNPs in non-coding regions of the human genome. Epigenetics and Chromatin. 2015.

67. Breen G, Li Q, Roth BL, O’Donnell P, Didriksen M, Dolmetsch R, et al. Translating genome-wide association findings into new therapeutics for psychiatry. Nat Neurosci. 2016.

68. Wu Z, Wang Y, Chen L. Network-based drug repositioning. Mol Biosyst. 2013.

69. Lotfi Shahreza M, Ghadiri N, Mousavi SR, Varshosaz J, Green JR. A review of network-based approaches to drug repositioning. Brief Bioinform. 2018.

70. Gillis J, Pavlidis P. ‘Guilt by association’ is the exception rather than the rule in gene networks. PLoS Comput Biol. 2012. 2012. https://doi.org/10.1371/journal.pcbi.1002444.

71. Mejía-Pedroza RA, Espinal-Enríquez J, Hernández-Lemus E. Pathway-based drug repositioning for breast cancer molecular subtypes. Front Pharmacol. 2018. 2018. https://doi.org/10.3389/fphar.2018.00905.

72. Martínez V, Navarro C, Cano C, Fajardo W, Blanco A. DrugNet: Network-based drug-disease prioritization by integrating heterogeneous data. Artif Intell Med. 2015. 2015. https://doi.org/10.1016/j.artmed.2014.11.003.

73. Li J, Lu Z. Pathway-based drug repositioning using causal inference. BMC Bioinformatics. 2013. 2013. https://doi.org/10.1186/1471-2105-14-S16-S3.

74. Wang W, Yang S, Zhang X, Li J. Drug repositioning by integrating target information through a heterogeneous network model. Bioinformatics. 2014. 2014. https://doi.org/10.1093/bioinformatics/btu403.

75. Luo H, Wang J, Li M, Luo J, Peng X, Wu FX, et al. Drug repositioning based on comprehensive similarity measures and Bi-Random walk algorithm. Bioinformatics, 2016.

76. Cheng T, Li Q, Wang Y, Bryant SH. Identifying compound-target associations by combining bioactivity profile similarity search and public databases mining. J Chem Inf Model. 2011. 2011. https://doi.org/10.1021/ci200192v.

77. Bellera CL, Balcazar DE, Vanrell MC, Casassa AF, Palestro PH, Gavernet L, et al. Computer-guided drug repurposing: Identification of trypanocidal activity of clofazimine, benidipine and saquinavir. Eur J Med Chem. 2015. 2015. https://doi.org/10.1016/j.ejmech.2015.01.065.

78. Maynard RL. The Merck Index: 12th edition 1996. Occup Environ Med. 1997. 1997. https://doi.org/10.1136/oem.54.4.288.

79. Tari LB, Patel JH. Systematic Drug Repurposing Through Text Mining. Methods Mol Biol. 2014. 2014. https://doi.org/10.1007/978-1-4939-0709-0_14.

80. Krallinger M, Erhardt RA-A, Valencia A. Text-mining approaches in molecular biology and biomedicine. Drug Discov Today. 2005;10:439–445.

81. Zheng S, Dharssi S, Wu M, Li J, Lu Z. Text Mining for Drug Discovery2019. p. 231–252.

82. Xue H, Li J, Xie H, Wang Y. Review of drug repositioning approaches and resources. Int J Biol Sci. 2018.

83. Li J, Zhu X, Chen JY. Building disease-specific drug-protein connectivity maps from molecular interaction networks and PubMed abstracts. PLoS Comput Biol. 2009. 2009. https://doi.org/10.1371/journal.pcbi.1000450.

84. WHO. ATC - Structure and principles. WHO Collab Cent Drug Stat Methodol. 2012.

85. Napolitano F, Zhao Y, Moreira VM, Tagliaferri R, Kere J, D’Amato M, et al. Drug repositioning: A machine-learning approach through data integration. J Cheminform. 2013. 2013. https://doi.org/10.1186/1758-2946-5-30.

86. Chen L, Zeng WM, Cai YD, Feng KY, Chou KC. Predicting anatomical therapeutic chemical (ATC) classification of drugs by integrating chemical-chemical interactions and similarities. PLoS One. 2012. 2012. https://doi.org/10.1371/journal.pone.0035254.

87. Liu Z, Guo F, Gu J, Wang Y, Li Y, Wang D, et al. Similarity-based prediction for Anatomical Therapeutic Chemical classification of drugs by integrating multiple data sources. Bioinformatics, 2015.

88. WHOCC - Structure and principles. https://www.whocc.no/atc/structure_and_principles/. Accessed 1 April 2020.

89. Edwards IR, Aronson JK. Adverse drug reactions: Definitions, diagnosis, and management. Lancet. 2000. 2000. https://doi.org/10.1016/S0140-6736(00)02799-9.

90. Kant A, Bilmen J, Hopkins PM. Adverse drug reactions. Pharmacol. Physiol. Anesth. Found. Clin. Appl., 2018.

91. Rohilla A, Yadav S. Adverse drug reactions: An Overview. Int J Pharmacol Res. 2013. 2013. https://doi.org/10.7439/IJPR.V3I1.41.

92. Wang Z, Clark NR, Ma’ayan A. Drug-induced adverse events prediction with the LINCS L1000 data. Bioinformatics. 2016. 2016. https://doi.org/10.1093/bioinformatics/btw168.

93. Kim S, Chen J, Cheng T, Gindulyte A, He J, He S, et al. PubChem 2019 update: Improved access to chemical data. Nucleic Acids Res. 2019. 2019. https://doi.org/10.1093/nar/gky1033.

94. Kuhn M, Letunic I, Jensen LJ, Bork P. The SIDER database of drugs and side effects. Nucleic Acids Res. 2016. 2016. https://doi.org/10.1093/nar/gkv1075.

95. Tatonetti NP, Ye PP, Daneshjou R, Altman RB. Data-driven prediction of drug effects and interactions. Sci Transl Med. 2012. 2012. https://doi.org/10.1126/scitranslmed.3003377.

96. Geurts P, Ernst D, Wehenkel L. Extremely randomized trees. Mach Learn. 2006. 2006. https://doi.org/10.1007/s10994-006-6226-1.

97. Mechanism matters. vol. 16. Nature Publishing Group; 2010.

98. Iorio F, Bosotti R, Scacheri E, Belcastro V, Mithbaokar P, Ferriero R, et al. Discovery of drug mode of action and drug repositioning from transcriptional responses. Proc Natl Acad Sci U S A. 2010. 2010. https://doi.org/10.1073/pnas.1000138107.

99. Madhukar NS, Khade PK, Huang L, Gayvert K, Galletti G, Stogniew M, et al. A Bayesian machine learning approach for drug target identification using diverse data types. Nat Commun. 2019. 2019. https://doi.org/10.1038/s41467-019-12928-6.

100. Friedman R. Drug resistance in cancer: Molecular evolution and compensatory proliferation. Oncotarget. 2016. 2016. https://doi.org/10.18632/oncotarget.7459.

101. Berger AH, Brooks AN, Wu X, Shrestha Y, Chouinard C, Piccioni F, et al. High-throughput Phenotyping of Lung Cancer Somatic Mutations. Cancer Cell. 2016. 2016. https://doi.org/10.1016/j.ccell.2016.06.022.

102. Pajouhesh H, Lenz GR. Medicinal chemical properties of successful central nervous system drugs. NeuroRx. 2005. 2005. https://doi.org/10.1602/neurorx.2.4.541.

103. Daina A, Michielin O, Zoete V. SwissADME: A free web tool to evaluate pharmacokinetics, drug-likeness and medicinal chemistry friendliness of small molecules. Sci Rep. 2017. 2017. https://doi.org/10.1038/srep42717.

104. Wawer MJ, Li K, Gustafsdottir SM, Ljosa V, Bodycombe NE, Marton MA, et al. Toward performance-diverse small-molecule libraries for cell-based phenotypic screening using multiplexed high-dimensional profiling. Proc Natl Acad Sci U S A. 2014. 2014. https://doi.org/10.1073/pnas.1410933111.

105. Sullivan CR, Mielnik CA, O’Donovan SM, Funk AJ, Bentea E, DePasquale EA, et al. Connectivity Analyses of Bioenergetic Changes in Schizophrenia: Identification of Novel Treatments. Mol Neurobiol. 2019. 2019. https://doi.org/10.1007/s12035-018-1390-4.

106. Altar CA, Jurata LW, Charles V, Lemire A, Liu P, Bukhman Y, et al. Deficient hippocampal neuron expression of proteasome, ubiquitin, and mitochondrial genes in multiple schizophrenia cohorts. Biol Psychiatry. 2005. 2005. https://doi.org/10.1016/j.biopsych.2005.03.031.

107. Stone WS, Faraone S V., Su J, Tarbox SI, Van Eerdewegh P, Tsuang MT. Evidence for linkage between regulatory enzymes in glycolysis and schizophrenia in a multiplex sample. Am J Med Genet. 2004. 2004. https://doi.org/10.1002/ajmg.b.20132.

108. Bang S, Jeong S, Choi N, Kim HN. Brain-on-a-chip: A history of development and future perspective. Biomicrofluidics. 2019;13:051301.

109. Xu H, Jiao Y, Qin S, Zhao W, Chu Q, Wu K. Organoid technology in disease modelling, drug development, personalized treatment and regeneration medicine. Exp Hematol Oncol. 2018;7:30.

110. Lopez-Munoz F, Alamo C. Monoaminergic Neurotransmission: The History of the Discovery of Antidepressants from 1950s Until Today. Curr Pharm Des. 2009. 2009. https://doi.org/10.2174/138161209788168001.

111. Wenthur CJ, Bennett MR, Lindsley CW. Classics in chemical neuroscience: Fluoxetine (Prozac). ACS Chem Neurosci. 2014.

112. Duman RS. The Dazzling Promise of Ketamine. Cerebrum. 2018;2018.

113. Alda M. Lithium in the treatment of bipolar disorder: pharmacology and pharmacogenetics. Mol Psychiatry. 2015;20:661–670.

114. Moret N, Clark NA, Hafner M, Wang Y, Lounkine E, Medvedovic M, et al. Cheminformatics Tools for Analyzing and Designing Optimized Small-Molecule Collections and Libraries. Cell Chem Biol. 2019. 2019. https://doi.org/10.1016/j.chembiol.2019.02.018.

115. Lin E, Lin CH, Lane HY. Precision psychiatry applications with pharmacogenomics: Artificial intelligence and machine learning approaches. Int J Mol Sci. 2020.

116. Baldwin DS, Kosky N. Off-label prescribing in psychiatric practice. Adv Psychiatr Treat. 2007.

117. Li YY, Jones SJM. Drug repositioning for personalized medicine. Genome Med. 2012.

118. Bentea E, Depasquale EAK, O’Donovan SM, Sullivan CR, Simmons M, Meador-Woodruff JH, et al. Kinase network dysregulation in a human induced pluripotent stem cell model of DISC1 schizophrenia. Mol Omi. 2019. 2019. https://doi.org/10.1039/c8mo00173a.

119. Spencer T, Biederman J, Heiligenstein J, Wilens T, Faries D, Prince J, et al. An open-label, dose-ranging study of atomoxetine in children with attention deficit hyperactivity disorder. J Child Adolesc Psychopharmacol. 2001. 2001. https://doi.org/10.1089/10445460152595577.

120. Schwartz J, Murrough JW, Iosifescu D V. Ketamine for treatment-resistant depression: recent developments and clinical applications: Table 1. Evid Based Ment Heal. 2016;19:35–38.

121. Ferry L, Johnston JA. Efficacy and safety of bupropion SR for smoking cessation: data from clinical trials and five years of postmarketing experience. Int J Clin Pract. 2003. 2003.

122. Corbett A, Ballard C. New and emerging treatments for Alzheimer’s disease. Expert Opin Emerg Drugs. 2012.

123. Wang J, Ho L, Chen L, Zhao Z, Zhao W, Qian X, et al. Valsartan lowers brain β-amyloid protein levels and improves spatial learning in a mouse model of Alzheimer disease. J Clin Invest. 2007. 2007. https://doi.org/10.1172/JCI31547.

124. Nickell JR, Grinevich VP, Siripurapu KB, Smith AM, Dwoskin LP. Potential therapeutic uses of mecamylamine and its stereoisomers. Pharmacol Biochem Behav. 2013.

125. Belanoff JK, Flores BH, Kalezhan M, Sund B, Schatzberg AF. Rapid reversal of psychotic depression using mifepristone. J Clin Psychopharmacol. 2001. 2001. https://doi.org/10.1097/00004714-200110000-00009.

126. Saraf G, Viswanath B, Hatti S, Malyala A, Benegal V. A comparison of baclofen and topiramate with acamprosate as anticraving agents: A naturalistic follow-up in a tertiary care de-addiction unit. Alcohol Clin Exp Res. 2012. 2012. https://doi.org/http://dx.doi.org/10.1111/j.1530-0277.2012.01803.x.

127. Gorsane MA, Kebir O, Hache G, Blecha L, Aubin HJ, Reynaud M, et al. Is baclofen a revolutionary medication in alcohol addiction management? Review and recent updates. Subst Abus. 2012. 2012. https://doi.org/10.1080/08897077.2012.663326.

128. Hayes JF, Lundin A, Wicks S, Lewis G, Wong ICK, Osborn DPJ, et al. Association of Hydroxylmethyl Glutaryl Coenzyme A Reductase Inhibitors, L-Type Calcium Channel Antagonists, and Biguanides with Rates of Psychiatric Hospitalization and Self-Harm in Individuals with Serious Mental Illness. JAMA Psychiatry. 2018. 2018. https://doi.org/10.1001/jamapsychiatry.2018.3907.

129. So H-CC, Chau CK-LL, Chiu W-TT, Ho K-SS, Lo C-PP, Yim SH-YY, et al. Analysis of genome-wide association data highlights candidates for drug repositioning in psychiatry. Nat Neurosci. 2017;20:1342–1349.

130. Kwok MK, Lin SL, Schooling CM. Re-thinking Alzheimer’s disease therapeutic targets using gene-based tests. EBioMedicine. 2018. 2018. https://doi.org/10.1016/j.ebiom.2018.10.001.

131. Zhang Y shuai, Li J dong, Yan C. An update on vinpocetine: New discoveries and clinical implications. Eur J Pharmacol. 2018.

132. Gaspar HA, Gerring Z, Hübel C, Middeldorp CM, Derks EM, Breen G. Using genetic drug-target networks to develop new drug hypotheses for major depressive disorder. Transl Psychiatry. 2019. 2019. https://doi.org/10.1038/s41398-019-0451-4.

133. Rodriguez-López J, Arrojo M, Paz E, Páramo M, Costas J. Identification of relevant hub genes for early intervention at gene coexpression modules with altered predicted expression in schizophrenia. Prog Neuro-Psychopharmacology Biol Psychiatry. 2020. 2020. https://doi.org/10.1016/j.pnpbp.2019.109815.

134. De Jong S, Vidler LR, Mokrab Y, Collier DA, Breen G. Gene-set analysis based on the pharmacological profiles of drugs to identify repurposing opportunities in schizophrenia. J Psychopharmacol. 2016. 2016. https://doi.org/10.1177/0269881116653109.

135. Lencz T, Malhotra AK. Targeting the schizophrenia genome: A fast track strategy from GWAS to clinic. Mol Psychiatry. 2015. 2015. https://doi.org/10.1038/mp.2015.28.

136. Wang S, Meng X, Wang Y, Liu Y, Xia J. HPO-Shuffle: An associated gene prioritization strategy and its application in drug repurposing for the treatment of canine epilepsy. Biosci Rep. 2019. 2019. https://doi.org/10.1042/BSR20191247.

